# Keeping an Eye on Plasticity Genes: Insulin/TOR Pathway Nodes Mediating Nutritional Plasticity of Eyes Within and Between Sex

**DOI:** 10.1101/2025.08.31.673361

**Authors:** Sarah Heffernan, Emma Etchells Foisy, Kadija Saleebaan, Rajendhran Rajakumar

## Abstract

The eyes of insects exhibit extreme morphological variation through changes in size and shape. While the model organism *Drosophila melanogaster* has been used to elucidate the underlying gene networks and cell-signalling pathways that regulate the patterning and growth of the eye, insight into the regulators of nutrient-dependent sizing and allometry within and between the sexes remains poorly understood. Using a screen approach of the well characterized and ancient Insulin/TOR pathway, we show that perturbations of different nodes of this pathway modulate the impact of nutritional variation on the sizing and scaling of the eye to varying degrees, and that this is further influenced by sex. When starved, wild-type flies decrease in eye size and body size proportionately in females, yet males change eye-to-body size disproportionately. Subsequently we used previously validated RNAi lines to knockdown each component of the Insulin/TOR pathway in an eye-specific manner to characterize their individual influence in modulating nutritional plasticity of eye sizing. Surprisingly, this resulted in a range of size and scaling variation, components modulating the plastic response on eye size mean, eye-to-body slope, intercept, as well as eye size variance, some effects ranging in magnitude from shutting off plasticity to amplifying, and some which were sex-limited. More generally, the morphospace of a trait can evolve within and across the sexes through modifications of plasticity genes.

## Introduction

Variation in the size of a morphological trait and how this variation can evolve has long intrigued biologists (Darwin, 1964; Hallgrímsson and Hall, 2011; Hamilton, 1961; Huxley et al., 1993; Thompson, 1992). Trait size can change in mean size, degree of variation, as well as its allometry, that is, the size of the trait in relation to body size (Stillwell et al., 2016). This size variation can be the product of both genetic variation and environmental variation acting on the developmental processes underlying the trait’s size (Frankino et al., 2019). Furthermore, sizing differences across the sexes can range from subtle to extreme (sexual size dimorphism) depending on the species (Fairbairn, 1997). Several cell-signalling pathways have been implicated in regulating trait sizing during development (Gokhale and Shingleton, 2015; Texada et al., 2020). The ability for the developmental growth of a trait to be influenced by several environmental factors, including nutrition and temperature, is known as developmental plasticity (Schlichting and Pigliucci, 1998; Sultan, 2015; West-Eberhard, 2003). Characterizing how these pathways are influenced by varying environments and both how genetic and epigenetic variation can contribute to gene-by-environment interactions is critical towards pinpointing the sources for size differences, and more generally, variation across traits (Lafuente and Beldade, 2019; Meaney, 2017; Vandermeulen and Cullen, 2022).

One of the most extensively investigated cell-signalling pathways underlying growth regulation is the Insulin/TOR pathway, which itself can sense nutritional variation and translate that into differential growth of traits and the organism as a whole (Oldham and Hafen, 2003; Saltiel and Kahn, 2001; Schmelzle and Hall, 2000). Examples implicating this pathway in regulating trait and organismal growth range from the body size of bees and dogs to the size of mouse hearts and beetle horns (Emlen et al., 2012; Hoopes et al., 2012; Patel et al., 2007; Shioi, 2000; Sutter et al., 2007). In flies the major components of the pathway include the insulin receptor (InR), the insulin receptor substrate (chico), the phosphoinositide-3 kinase (PI3K), the protein kinase B (Akt), the phosphatase and tensin homolog (PTEN), the transcription factor Forkhead box-O (FOXO), *Drosophila melanogaster* homolog of Target of rapamycin (dTOR), the tuberous sclerosis complex proteins 1 and 2 (TSC1 and gigas/TSC2), the small GTPase Ras homolog enriched in brain (Rheb), the regulatory associated protein of the TOR complex (Raptor), the 4-E binding protein (Thor), and the S6-kinase (S6k) (Edgar, 2006; Grewal, 2009). Classic model organism genetic work has been done on the Insulin/TOR pathway in both the mouse and *Drosophila melanogaster* (Garofalo, 2002; Oldham and Hafen, 2003; Wullschleger et al., 2006). Furthermore, there are several examples demonstrating how this pathway can influence allometry. Tang et al., functionally scanned the Insulin/TOR pathway for factors capable of influencing the degree of plasticity of sizing in *D. melanogaster* and found that FOXO limits the nutritional plasticity of genitalia compared to the nutritional plasticity of the wing (Tang et al., 2011).

Extensive genetic research in *D. melanogaster* has been done using the eye as a phenotypic readout of growth and sizing to characterize the role of Insulin/TOR pathway components in growth regulation and size variation. For example, Gain-of-function (GOF) of *Pten* leads to eye reduction and loss-of-function (LOF) leads to overproliferation in the eye (Gao et al., 2000; Huang et al., 1999), GOF of the insulin receptor (*InR*) and insulin receptor substrate (*chico*) leads to eye size increase and LOF reduces eye size (Böhni et al., 1999; Brogiolo et al., 2001), and GOF of *Tsc1* and *Tsc2* lead to an increase in eye size in contrast to LOF leading to eye reduction (Gao and Pan, 2001; Tapon et al., 2001). Based on this extensive characterization of Insulin/TOR signalling and eye size regulation, the goal of this study is to use the *Drosophila melanogaster* eye as a system to investigate the influence of individual components of the Insulin/TOR pathway on nutrient-dependent size, size variation, and allometry variation within and between sex. To do this, we first exposed developing wildtype flies to varying nutritional conditions. We then characterized the effect of these conditions on mean eye size, eye size variation, as well as eye-body allometry. Next, we combined this nutritional experimental setup with eye-specific molecular genetic perturbations of all the major Insulin/TOR components in both males and females and characterized the effect of each pathway component on eye size and allometry in terms of mean size, slope, intercept, and degree of variation. This screen took advantage of previously validated RNAi lines that have been extensively characterized at the cellular, molecular, and phenotypic levels in the context of the growth of eyes as well as other organs in *D. melanogaster* (Birnbaum et al., 2021; Bradshaw et al., 2024; Lee et al., 2026; Millington et al., 2021; Myers et al., 2022; Nechipurenko and Broihier, 2012; Pletcher et al., 2019; Pranoto et al., 2023; Recasens-Alvarez et al., 2021; Rojas Villa et al., 2019; Trivedi et al., 2020; Zhao et al., 2025). Altogether, by leveraging the genetic model *D. melanogaster* and eye variation to study the interplay of nutritional plasticity and sexual dimorphism in the context of sizing we were able to identify unisex and sex-limited plasticity genes capable of influencing the environmental response of an individual’s developing eye and therefore facilitating potentially adaptive sizing and scaling possibilities.

## RESULTS

### Nutritional Plasticity and Eye sizing-scaling

To address the impact of individual components of the Insulin/TOR pathway on the effects of nutritional plasticity on sizing and allometric eye-to-body variation in males and females, we first characterized the eye size, allometry, and variance of eye size in different nutritional conditions. Wildtype *D. melanogaster* adults were placed in vials and larvae were isolated at the third and final instar and placed into either a fed or starved condition. After the feeding condition exposure, developing individuals were moved back to vials with food and left to reach adulthood. For comparisons, to account for a potential interaction of sex and nutritional condition, we did a two-way ANOVA for mean eye size, two-way ANCOVA for allometry (slopes and y-intercepts), and a two-way Levene’s test for variance. As compared to the fed condition, starved females had a significant decrease in body size (data not shown; *P*<0.0001), and eyes size (eye width) (Fig. 1A; *P<*0.0001), while their eye allometry remained the same with no significant change in slope or intercept (Fig. 1B). Similarly, as compared to the fed condition, starved males had a significant decrease in body size and eye size, yet there was a significant increase in slope (Fig. 1B). Therefore, while starvation causes a change in both body size and eye size across males and females, it does so proportionally in females, yet disproportionately in males. Next, we wanted to determine whether there was a change in variance of eye size within each of the nutritional conditions and/or a change in variance across conditions. As compared to fed conditions, starvation significantly increased the variance in eye size for females (1.87-fold; Fig. 1C; *P=0.0114*). Finally, we found there was no interaction between sex and nutrition for mean eye size, and variance, but there was for allometry (Fig. 1B; *P=*0.0125) for these wildtype flies. Altogether, nutritional conditions have both similar and sex-specific effects on eye size, allometry, and variance (Table S1).

**Fig. 1.**
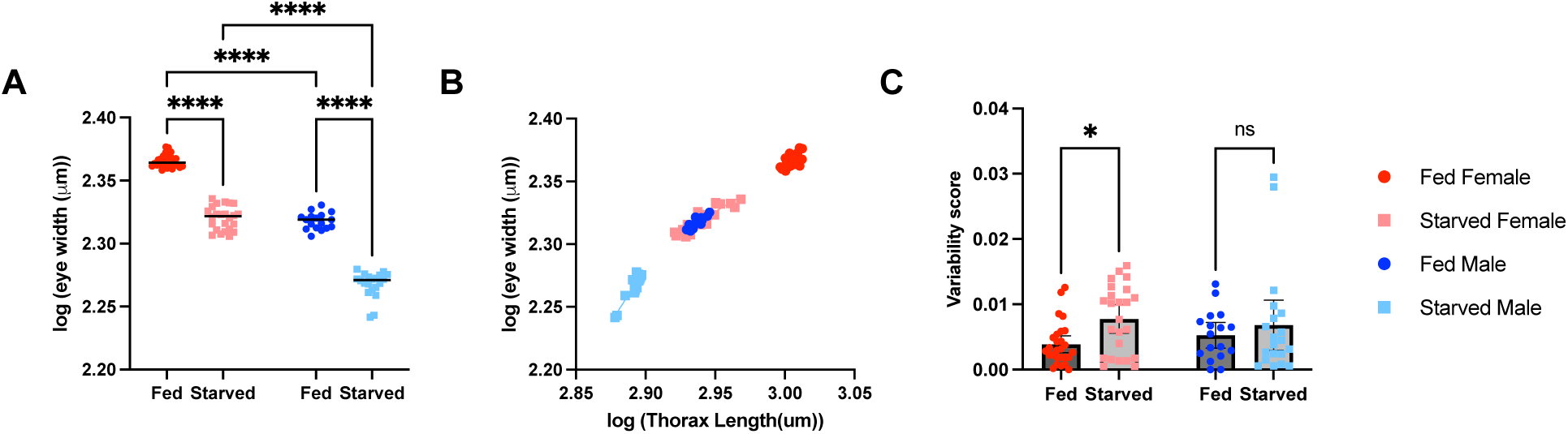
Effect of starvation on eye size in wildtype Oregon-R adult females and males. (A) Eye width across males and females under different developmental nutritional conditions (fed vs starved). Statistical significance for two-way ANOVA comparisons: ****P<0.0001. (B) The scaling relationship between eye width against body size in fed vs starved females and males. (C) Absolute deviation from median log(eye width) for fed and starved females and males (means ± 95% CI). Statistical significance for two-way ANOVA comparisons: *P<0.05. Female fed n=27(dark red circles), female starved n=24 (light red squares), male fed n=17(dark blue circles), male starved n=20 (light blue squares). Variability score = |log(eye width) - median log(eye width)| which is used for a two-way Levene’s Test.

### Insulin-TOR components as modulators of eye nutritional plasticity

To determine the role that the Insulin/TOR pathway plays in modulating the degree of nutritional plasticity of eye size, allometry, and variation across males and females we took a network approach. We functionally manipulated individual components of the Insulin/TOR pathway during the development of the eye imaginal discs, the population of cells that pattern and grow into the eye, in an eye-specific manner in both nutritional conditions using the Gal4-UAS system (Brand and Perrimon, 1993). In addition to the intracellular components of the Insulin/TOR pathway, we further included the highly conserved ligand Insulin-like peptide 2 (Dilp2) and the insulin receptor decoy, Imaginal morphogenesis protein-Late 2 (ImpL2) to investigate their role in eye nutrition plasticity as they are both expressed within imaginal tissue in parallel to their interorgan signalling activity (Brogiolo et al., 2001; Klebes et al., 2002).To do this, we used an eye-specific Gal4 line and crossed this to Insulin/TOR component-specific UAS RNAi lines (as well as a UAS-control ‘empty’ line) and repeated the same nutritional conditions as was done with wildtype. For comparisons, to account for a potential interaction of sex and nutritional condition, we did a two-way ANOVA for mean eye size, two-way ANCOVA for allometry (slopes and y-intercepts), and a two-way Levene’s test for variance. For the negative control, all sizing and allometry patterns when comparing sex and nutritional conditions were consistent with wildtype (Fig. 2A, B, Fig. 3, Table S1). Through measurements of eye width, there was a reduction in eye size in starved individuals as compared to fed for all Insulin/TOR perturbations except for *foxo* and *Rheb*, which both exhibited a non-significant difference in males and females (Fig. 2A,B, Fig. 3, Table S1). The lack of change in eye size after perturbing *foxo* and *Rheb* (males and females) is in stark contrast to the change in body size in the two nutritional conditions (Fig. S1).

**Fig. 2.**
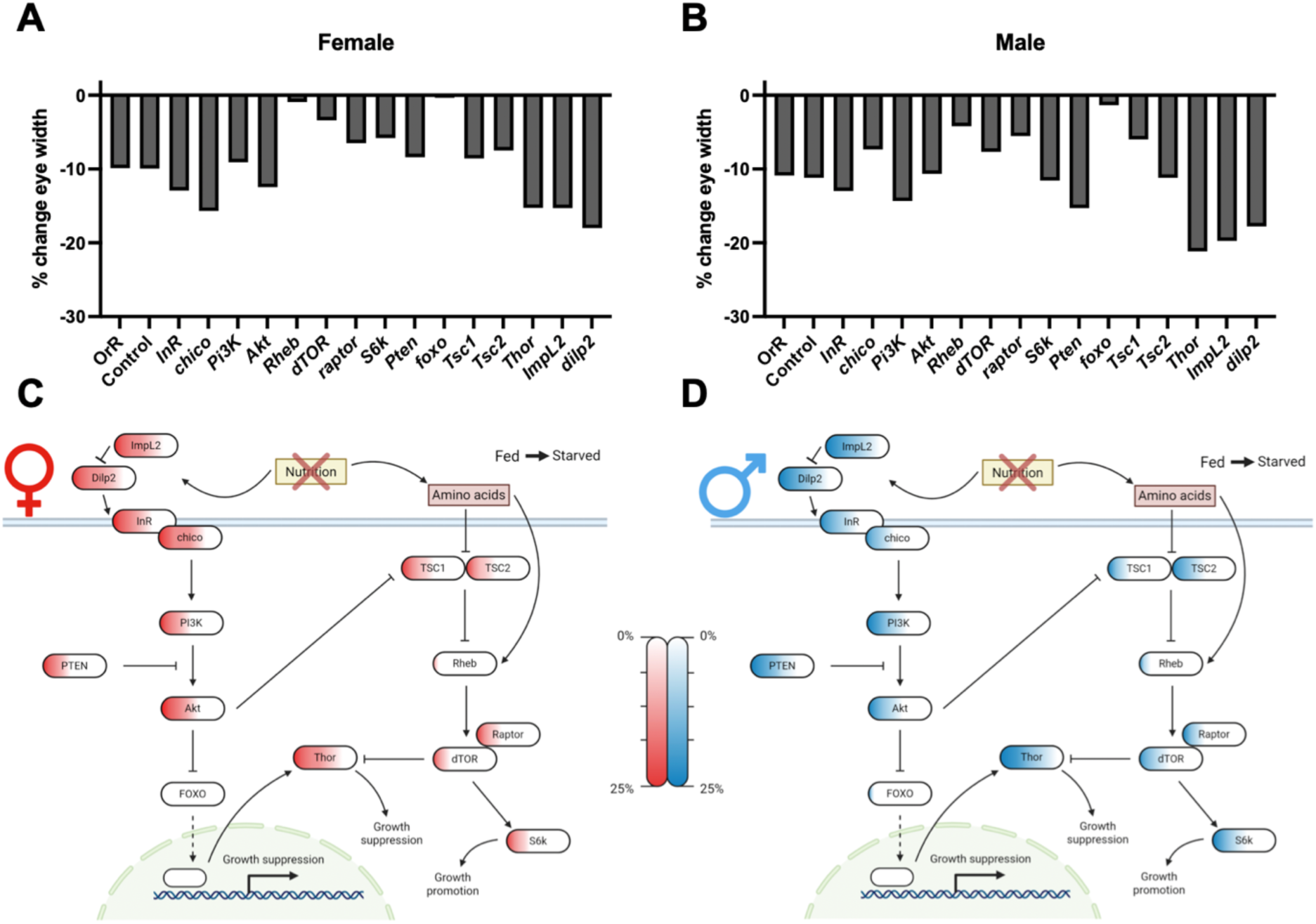
Effect of eye-specific Insulin/TOR pathway knockdown on percent change in eye size under different nutritional conditions. Percent change in eye width between fed and starved (A) females and (B) males. Interactions of the Insulin/TOR genes and their corresponding percent changes in eye width between fed and starved (C) females and (D) males when knocked down in the eye. Red/blue gradient represents percentage change from a scale of 0 to 25% for the respective gene. The more the red/blue colour gradient fills the gene box, the higher the percent change. Created in BioRender.

**Fig. 3.**
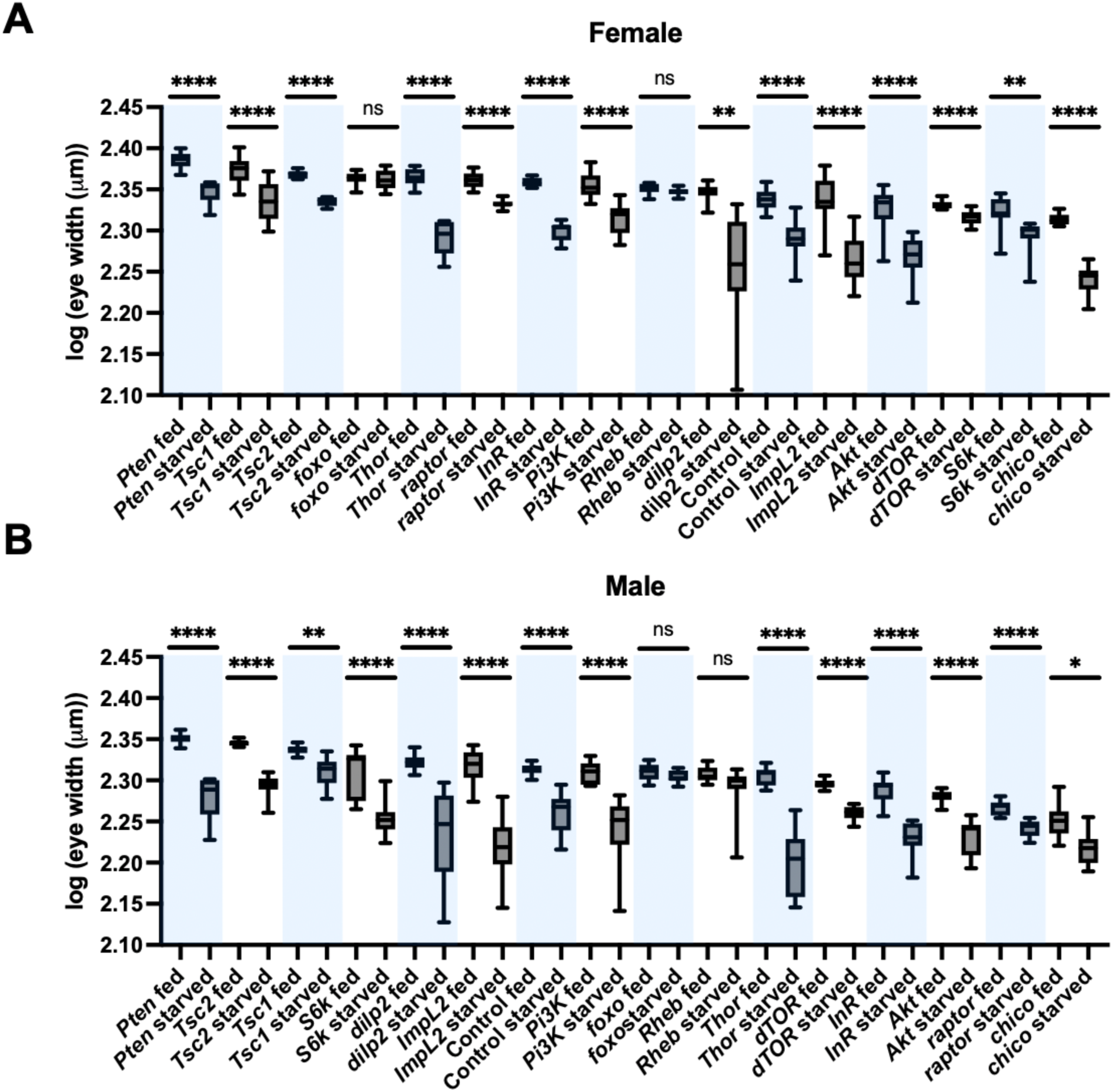
Comparison of absolute eye size of fed and starved Insulin/TOR knockdown females and males. (A) Absolute eye size (log(eye width (µm))) between fed and starved females when components of the Insulin/TOR pathway are individually knocked down in the eye. (B) Absolute eye size (log(eye width (µm))) between fed and starved males when components of the Insulin/TOR pathway are individually knocked down in the eye. Box of boxplots is defined by the 25^th^ and 75^th^ percentile with line at the median. Whiskers indicate min and max points. Statistical significance based on Holm-Bonferroni adjusted *P*-values from two-way ANOVA: \**P*<0.05, \*\**P*<0.01, \*\*\*\**P*<0.0001.

Furthermore, we found an interaction between sex and nutritional condition for mean eye size change for several genes. Specifically, there was an interaction for *S6k* (*P*=0.0487), *Thor* (*P*=0.0344), *dTOR* (*P*=0.0002), *Pten* (*P*=0.0008), *Tsc2* (*P*=0.0013) that exhibited a greater change in mean eyes size in males compared to females while there was an interaction for *chico* (*P*=0.0012), where there was a greater change in mean eye size in females.

We next characterized how modulating Insulin/TOR components impact allometric changes of eye variation in response to nutritional conditions in males and females. For starved females as compared to fed, there was no significant difference in slopes for *InR*, *chico*, *Akt*, *Rheb*, *S6k, foxo, Tsc1*, *Tsc2*,*Thor, ImpL2,* and *dilp2* perturbations in the eye (Table S1, Fig. 4A-D, Table S2). However, knockdown of *foxo*, *Thor*, *InR*, and *chico* led to a significant difference in the y-intercept in starved compared to fed females (Table S3). Following knockdown of *Pi3K*, *dTOR*, *raptor*, and *Pten,* starved female flies had a significantly different slope (Table S1, Fig. 4A-D, Table S2). Specifically, *Pten* had an increase in slope in starved females compared to fed while, *Pi3K, dTOR,* and *raptor* had a decrease in slope in starved females compared to fed. In starved males (as compared to fed males), there was no significant difference when *InR, Tsc1*, *Tsc2*, *Thor*, *foxo*, *chico*, *Akt*, *Rheb*, *dTOR*, *S6k*, and *ImpL2* are knocked down in the eye (Table S1, Fig. 4E-H, Table S2). However, for *InR*, *Akt*, *dTOR*, *Thor*, and *Tsc2* there was a significant difference in the y-intercept in starved compared to fed males (Table S1 Fig. 4E-H, Table S3). Following knockdown of *Pi3K*, *raptor*, *Pten,* and *dilp2* starved male flies had a significantly different slope (Table S1, Fig. 4E-H, Table S2). Specifically, *Pi3K*, *Pten*, and *dilp2* had an increase in slope in starved males compared to fed, while *raptor* had a decrease in slope in starved males compared to fed. Furthermore, we found an interaction between sex and nutrition for *Pi3K* (*P*=0.0047) where the slope decreased in females between fed and starved (*P=*0.0337), while the slope increased in males between fed and starved (*P=*0.0069).

**Fig. 4.**
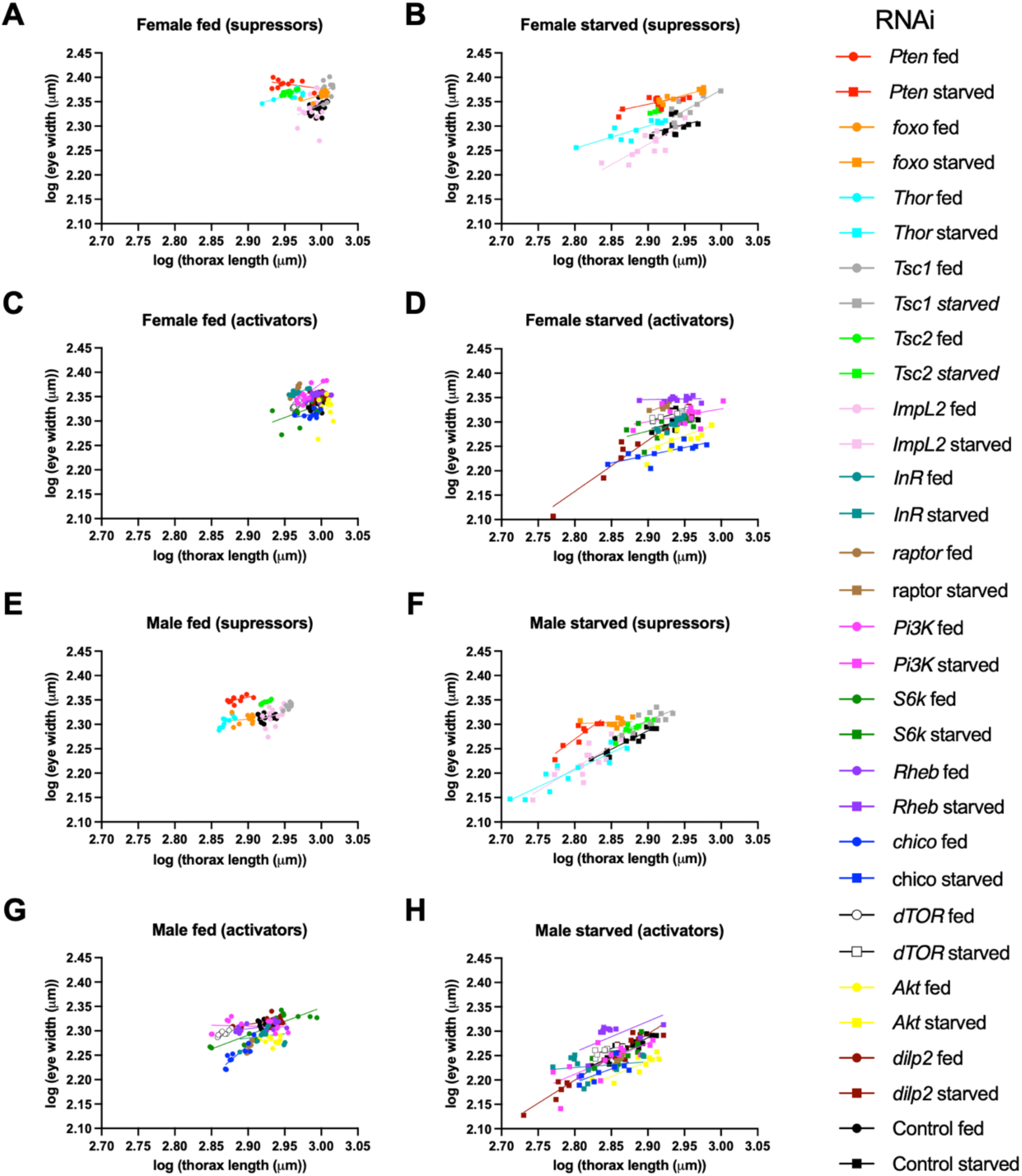
Allometric relationship between fed and starved Insulin/TOR knockdown females and males. Log-log plot of eye width-to-body allometry of (A) female fed and (B) female starved for Insulin/TOR pathway components which suppress growth. Log-log plot of eye width-to-body allometry of (C) female fed and (D) female starved for Insulin/TOR pathway components which activate growth. Log-log plot of eye width-to-body allometry of (E) male fed and (F) male starved for Insulin/TOR pathway components which suppress growth. Log-log plot of eye width-to-body allometry of (G) male fed and (H) male starved for Insulin/TOR pathway components which activate growth. Suppressors: *Pten*, foxo, *Thor*, *Tsc1*, *Tsc2*, *ImpL2.* Activators: *InR*,, *raptor*, *Pi3K*, *S6k*, *Rheb*, *chico*, d*TOR*, *Akt*, *dilp2*. Control is present in all plots. Refer to Tables S2 and S3. for slopes and y-intercepts.

Next, we wanted to determine whether perturbing components of the Insulin/TOR pathway would lead to a change in variance across nutritional conditions in males and females and how the different components ranked in terms of relative change in variance. For variance, *dilp2* increased in variance for both males (*P=*0.0004) and females (*P=*0.0003), while *Thor* (*P=*0.0004), *Pten* (*P=*0.0092), and *Tsc2* (*P=*0.0173) increased in variance for males and *raptor* (*P=*0.0399) decreased in variance in females (Fig. S2). Furthermore, we found an interaction between sex and nutrition for *Tsc1* (*P=*0.0443) (Fig. S2). Next, by comparing variances across Insulin/TOR perturbations by ranking the level of variance of each (for both sexes and both nutritional conditions), we wanted to better appreciate how nodes differed on a relative level. To do this, we determined the coefficient of variance for fed females, starved females, fed males, and starved males for each of the Insulin/TOR component perturbations. Based on the rank, from highest coefficient of variation to least, the following are the top 3 for the 4 conditions: (1) Fed females: *ImpL2*, *Tsc1*, and *Akt*; (2) starved females: *dilp2*, *ImpL2*, and *Tsc1;* (3) fed males: *S6k*, *chico*, *ImpL2*; and (4) starved males: *dilp2*,*Thor*, *Pi3K* (Table S1).When comparing fed to starved within each sex, we find that *raptor* RNAi leads to the biggest decrease in variation in females (rank 7 in fed females to 14 in starved females, Table S1), while *foxo* is the equivalent in males (rank 7 in fed males to 16 in starved males, Table S1). In contrast, *dilp2* RNAi leads to the biggest increase in variation in both females (rank 6 in fed females to rank 1 in starved females, Table S1), and males (rank of 9 in fed males to rank of 1 in starved males, Table S1). Collectively, depending on the sex, the nutritional conditions, and the interplay between the two, different components of the Insulin/TOR pathway can generate both a decrease and increase in eye size, allometry, and eye size variance.

## DISCUSSION

### Eye Sizing and Scaling: Model Organism to Model Clade

Based on our nutritional manipulations and eye-specific molecular perturbation experiments, we have used the *D. melanogaster* eye as a system to screen the ancient nutrient-sensing Insulin/TOR pathway for modifiers of size, allometry, and magnitude of variation in the context of nutritional plasticity. We have identified plasticity genes capable of modulating how the developing eye responds to an environmental condition as well as how the eye changes across environmental conditions (Table S1, Fig. 5). These changes come in the form of mean eye size, allometric changes in the slope or intercept as well as changes in trait variance. Furthermore, we have shown that these modifications can occur in a similar manner across sexes, yet they can also unfold in a sex-limited manner (Table S1, Fig. 5).

**Fig. 5.**
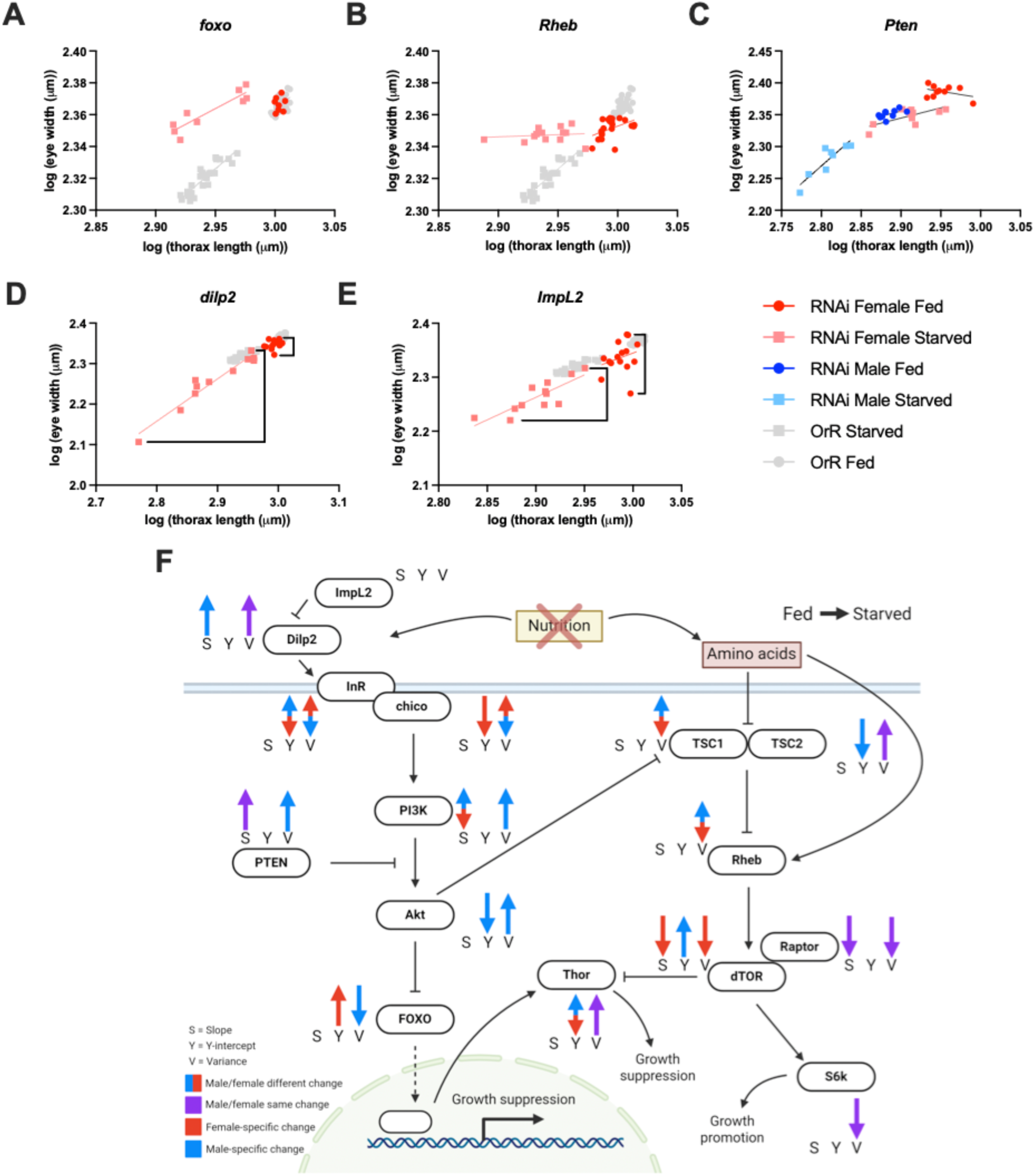
Examples of plasticity gene types. Across the Insulin/TOR pathway, components are capable of influencing sizing and scaling in different directions and in both sexes as well as in sex-limited ways. (A),(B) wild-type in grey, RNAi phenotypes in red, fed (circles) and starved (squares). For example: (A) *foxo* knockdown has negligible effect on mean eye size for fed females compared to wildtype. In contrast, *foxo* knockdown in a starved condition generates females with the same eye size as fed females yet this is much bigger than wildtype starved females. (B) *Rheb* knockdown mean eye size in starved females is also the same as *Rheb* knockdown in fed females, yet this similarity emerges in a different way than that found with *foxo*: fed *Rheb* knockdown female eyes are smaller than fed wildtype female eyes, yet starved *Rheb* knockdown female eyes are larger than wildtype. (C) *Pten* knockdown resulted in a decrease in mean eye size and an increase in slope for both starved females (red) and starved males (blue) compared to their fed counterparts. (D), *dilp2* knockdown resulted in an increase in variance of starved females while fed females did not change variance, whille (E) *Impl2* knockdown resulted in an increase in variance in both starved and fed conditions in a context-independent manner. (F) Summary of the Insulin-TOR pathway components’ impact on slope, y-intercept, and variance in males and females in the context of fed compared to starved nutrition conditions. Significant effects are represented as male (blue), female (red), both sexes (purple), up arrows are increase and down arrows are decrease. S is slope, Y is y-intercept, and V is variance. Created in BioRender.

Eye size is regulated by a highly conserved gene regulatory network (GRN) and several other cell-signalling pathways underlying patterning and growth of the developing eye (Casares and Almudi, 2016; Casares and McGregor, 2021; Kumar, 2010; Kumar, 2011). For instance, *eyeless*/*Pax-6* has long been known as a master regulator of eye development (Halder, Callaerts, and Gehring, 1995). Cis-regulatory evolution of *eyeless* across *Drosophila* species, as well as a single nucleotide polymorphism of a binding site in the *eyeless* cis-regulatory region for the transcription factor *cut* at the population level, has been implicated in the evolution of eye sizing across *Drosophila* (Ramaekers et al., 2019). How Insulin/TOR and other growth pathways integrate with patterning and specification components of the eye GRN at the population level and across *Drosophila* should be further explored. At the level of functional morphology, differences in overall eye size and the number and sizes of ommatidia in *Drosophila* species have functional implications on spatial acuity, temporal resolution, and contrast sensitivity (Buffry et al., 2024) and at the ecological-evolutionary level, the evolutionary transitions in eye size variation, and its sensory and life-history relevance has been traced across the *Drosophila* genus (Keesey et al., 2019). The contributing roles of the eye GRN and the Insulin/TOR pathway in regulating eye size can be situated within a larger framework that considers functional morphology, ecology, and evolutionary transitions. This model clade approach (Sanger and Rajakumar, 2019) leads to the following exciting question: how does the nutrient-sensing Insulin/TOR pathway integrate with the eye GRN to regulate cell size, proliferation, and eye-to-body allometry in varying nutritional contexts and how does this interaction contribute to the evolution of adaptive eye size variation and allometry across *Drosophila*?

### Insulin/TOR signalling from Established Models to Emerging Models

The Insulin/TOR pathway has been implicated in the development and evolution of the size of several developmentally plastic appendages and morphs. In *Apis mellifera*, both *TOR* and *chico* knockdown blocks the development of queen-destined larvae that are provided a queen diet, generating worker-like smaller sizes and reproductive tissues (Mutti et al., 2011; Patel et al., 2007; Wolschin et al., 2011). Furthermore, *Pten*, the inhibitor of the insulin branch of the Insulin/TOR pathway increases in expression when *A. mellifera* queen-destined larvae are given an inferior worker-diet (Wheeler et al., 2006). In rhinoceros beetles, well fed male larvae grow a massive head horn while malnourished males do not, and knockdown of the *insulin receptor* of male larvae fed optimal diets leads to the production of a significant reduction in the horn (Emlen et al., 2012). In *Onthophagus* horned beetles, *foxo* knockdown leads to an increase in horn and copulatory organ size (Snell-Rood and Moczek, 2012) and a change in the sigmoidal-shaped allometry of horn-to-body size between hornless and horned males (Casasa and Moczek, 2018). In an evolutionary context, in ants, when looking across 163 ant species reflecting all major ant groups, the two *insulin receptor* ant paralogs *InR1* and *InR2* have undergone selection across lineages with high levels of worker-queen size dimorphism and worker polymorphism while *chico* has undergone selection across lineages with high levels of worker-queen size dimorphism as well as relaxed selection for ommatidia number of the eye and *Pten* for queen-worker size dimorphism (Vizueta et al., 2025). Alongside the approach summarized above that leverages the model *D. melanogaster* for mechanistic resolution in tandem with the model clade *Drosophila* approach for evolutionary insight, these emerging model organisms ranging from bees to beetles altogether can offer a grander understanding of how the Insulin/TOR pathway may contribute to the development and evolution of eyes and traits more generally.

### Eyes Illuminate Plasticity Genes

A plasticity gene is a gene that can respond to an environmental cue triggering a specific series of developmental events, and allelic variation of a plasticity gene can lead to variation in the sensitivity of the plastic response (Pigliucci, 2001; Schlichting and Pigliucci, 1993; Via et al., 1995). Since the Insulin/TOR pathway is capable of sensing environmental variation, especially nutritional variation, its components have the capacity to be plasticity genes. By perturbing each of their functions in a tissue-specific manner, we have uncovered several plasticity genes, but more importantly, we have characterized the specific sizing and allometric parameters that can be modulated in the context of GxE interactions, the G through Insulin/TOR component perturbations and the E through nutritional conditions. The parameters that we were able to assess included trait mean, variance, and the slope and intercept of eye-to-body allometry (Fig. 5F). For example, *foxo* RNAi influenced the outcomes of fed females versus starved females by blocking the expected decrease in eye size that usually accompanies the decrease in body size in starved versus fed conditions (Fig. 2A,B, Fig. 3, Fig. 5A). While *foxo* RNAi did not affect eye size in fed females compared to wildtype, *foxo* prevented eye size reduction of starved developing individuals (Fig. 5A). *foxo* RNAi also led to a decrease in y-intercept in starved individuals compared to fed, which does not happen in wildtype nutritional conditions where females isometrically decrease in size (Fig. 1C). Furthermore, *foxo* fed and starved females both had higher variance than what is found in the wildtype (Table S1). It has been previously shown that wings are more nutritionally plastic than genitalia in *D.* melanogaster and a critical reason for this is that *foxo* plays a more significant plasticity role in wings than in the genitalia (Tang et al., 2011). Specifically, *foxo* overexpression leads to a greater reduction in *D. melanogaster* wing size than genitalia size and that increasing *foxo* in developing genitalia can lead to increased plasticity in the context of genitalia-to-body allometry (Tang et al., 2011). This indicates that *foxo* has the capacity to influence tissue-specific plastic responses to nutrition. In the context of the eye, increased expression of *foxo* leads to reduced eye size, similar to that found with decreased insulin signalling (Jünger et al., 2003; Kramer et al., 2003; Puig et al., 2003). Surprisingly, when insulin signalling is lowered (*chico* mutation), and simultaneously *foxo* is mutated, the *foxo* loss of function blocks the expected eye size reduction (Jünger et al., 2003). This is reminiscent of our findings that eye-specific loss of *foxo* prevents the eye from reducing in size in a coordinated manner with the body. Therefore, without *foxo*, the eye cannot properly respond to change in nutritional conditions. Similarly in the case of the *Onthophagus* horned beetles, *foxo* loss-of-function dampened the sigmoidal switch-like effect that nutritional conditions have (typically big horns in optimal nutrition, little-to-no horns in suboptimal)(Casasa and Moczek, 2018). These results in the horned beetle are consistent with ours in that, as a beetle decreases in body size lower than a precise threshold, usually there is a dramatic decrease in horn size (hyperplastic) yet when *foxo* is knocked down, this drop in horn size is dampened (Casasa and Moczek, 2018). In the case of *Onthophagus* beetles, this change in the structure of horn-to-body allometry and variation can have fundamental implications in the success of these beetles as the ability to precisely produce alternative horned and hornless males is essential to provide alternative reproductive strategies that these individuals employ in a morph-specific manner. Altogether, *foxo* is a nutrition-dependent plasticity gene that may be providing raw materials for natural and sexual selection.

In addition to *foxo*, our screen has found another plasticity gene that at first appears to block the effects of nutritional conditions similar to *foxo*. The percent change in size of starved versus fed females and males after the knockdown of *Rheb* leads to a near-zero percent change in eye size similar to that found with *foxo* (Fig. 2A,B, Fig. 3). Similarly to *foxo* RNAi that led to eye size being the same between the two nutritional conditions, *Rheb* starved and fed females and males also had the same eye size (Fig. 2), Fig. 3). Surprisingly, in females this is due to *Rheb* RNAi causing a decrease in eye size of fed females compared to wildtype and an increase in eye size in starved females compared to wildtype, which is different than what *foxo* RNAi generates (Fig. 5A and B). A mechanistic explanation could be found in past observations of *Rheb* and eye development in flies. Specifically, *Rheb* mutants inhibit eye growth, *Rheb* overexpression increases eye size (Saucedo et al., 2003; Stocker et al., 2003), and more importantly, starvation induces a rapid and sustained increase in *Rheb* expression and starved cells expressing *Rheb* are able to grow and activate *S6k* to regulate growth (Saucedo et al., 2003). Therefore, it is possible that our eye-specific *Rheb* RNAi in females in fed conditions led to the expected decrease in eye size and our increase in eye size in the starved conditions could be due to a rescue of *Rheb* expression that is caused by starvation conditions. This nuance of *Rheb* being a positive regulator of growth, but also capable of being upregulated in starved conditions may explain the resulting no percent change in eye size between *Rheb* RNAi females in both nutritional conditions. For males, while *Rheb* RNAi males similarly exhibit no percent change in both nutritional conditions, this is the outcome of fed *Rheb* RNAi males dramatically reducing their eye size equivalent to wildtype starved males, yet starved *Rheb* RNAi males remaining the same size as wildtype starved males (Fig. 4 G,H). Furthermore, *Rheb* knockdown in males did not lead to a change in eye-to-body allometry, which does occur in wildtype males when comparing nutritional conditions (Table S1). In the context of variance, when comparing fed versus starved conditions, the coefficient of variance went down with female *Rheb* RNAi flies, however males increased variance (Table S1). Depending on how an organism evolves its ability (increasing or dampening its sensitivity) to alter its body size in relation to nutritional conditions, this duality of *Rheb* may lead to complex changes in eye-to-body sizing, allometry, and variance.

A plasticity gene of the Insulin/TOR pathway capable of influencing eye size variation and the ability to dramatically change the magnitude of variation that nutritional variation can cause in a context-dependent manner is *dilp2*. While *dilp2* RNAi did not influence eye size variance in the fed condition, knockdown of *dilp2* led to a dramatic increase in eye size variance in starved individuals (Fig. 5D, fig.S2). In stark contrast, while *Impl2* RNAi caused a significant decrease in eye size similar to *dilp2* (Fig. 2, Fig. 3), it caused a radical increase in variance for both fed and starved conditions, far beyond the variation seen in wildtype developing individuals subjected to the two nutritional conditions (Fig. 5E). Therefore, while *dilp2* regulates variance in a context-dependent manner (fed-only), *Impl2* regulates variance in a context-independent manner (fed and starved). In particular, the change in variance observed for *dilp2* RNAi happened without perturbing the natural allometry of males and females, suggesting that variance is a parameter that plasticity genes can modulate independent of allometry (Table S1). *foxo*, *Rheb*, *dilp2* and *Impl2* are of the most unique factors we found thus far, nevertheless when looking across the Insulin/TOR pathway, we are able to detect plasticity genes that are able to modulate nutritional plasticity in different combinations of mean size, allometry (slope or intercept), and variance within a sex (Fig. 2C,D, Fig. 5, Fig. 6).

**Fig. 6.**
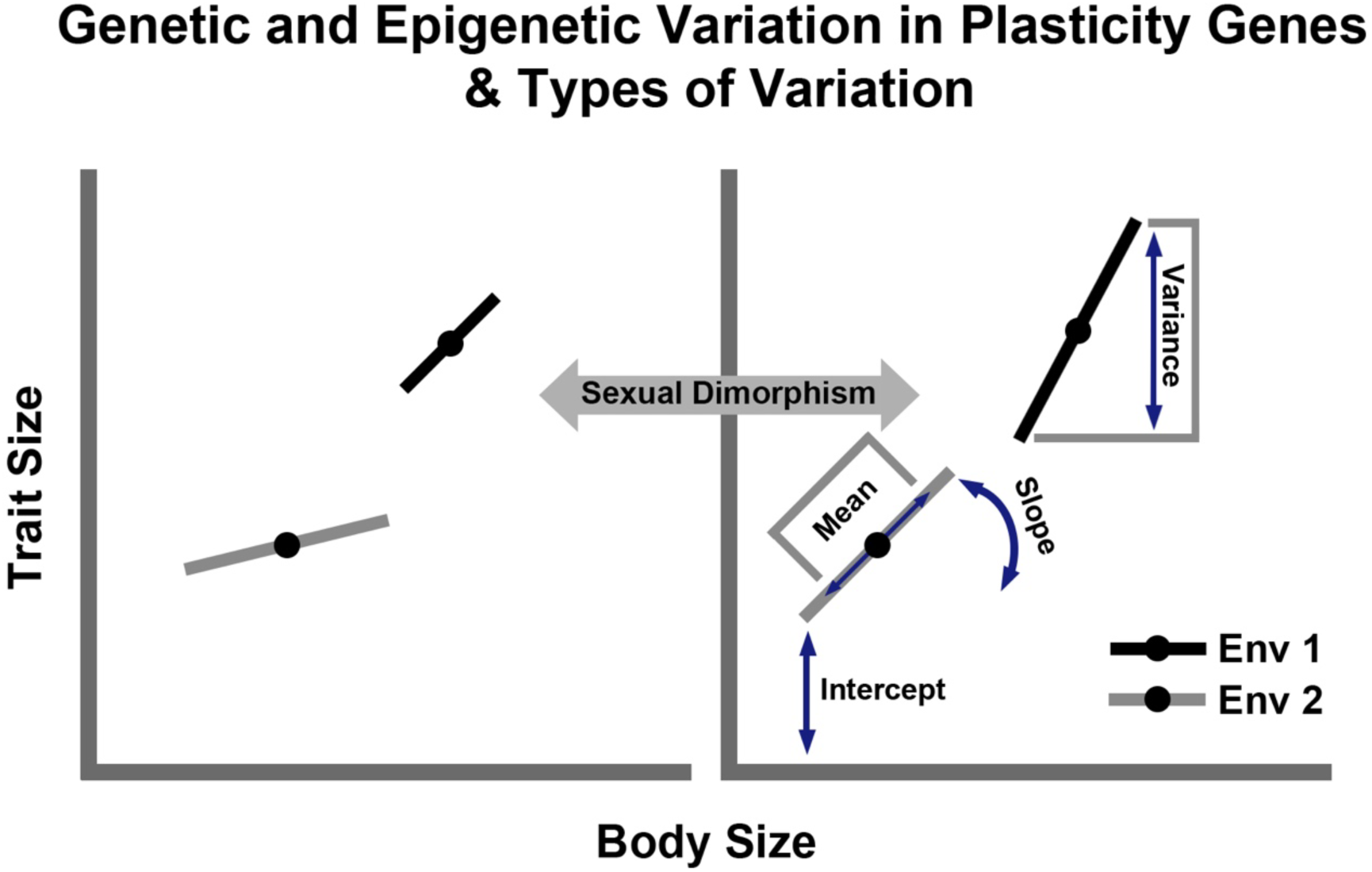
Types of plasticity genes Underlying Types of sizing and allometric variation. The Insulin/TOR pathway includes several critical anciently conserved components. When the functioning of different components is perturbed, different forms of phenotypic variation resulting from environment change emerge. In the context of eye size, 5 types of plasticity genes have been uncovered that impact: (1) the slope; (2) the intercept; (3) the mean independent of allometric change; (4) the degree of variance; (5) types 1-4 (slope, intercept, mean, or variance), but in a sex-limited manner. Changes in plasticity genes can emerge from both genetic and epigenetic sources.

### Sexual Dimorphism of Eye Plasticity Genes

Beyond our findings of Insulin/TOR plasticity genes having the ability to modulate size and allometry of eyes within a sex, we were also able to detect fascinating sexually dimorphic sex-limited roles that Insulin/TOR pathway plasticity genes provide. The origin and significance of phenotypic variation within and between sexes has long intrigued biologists (Andersson, 1994; Darwin, 1872; Miller and Svensson, 2014). Extensive investigations have been done to understand the influence of ecology (Cornwallis and Uller, 2010; Maan and Seehausen, 2011; Shine, 1989; Slatkin, 1984) and developmental processes (Badyaev, 2002; Hopkins and Kopp, 2021; Williams and Carroll, 2009) on the evolution of intra- and intersexual differences. An exciting avenue of current research is evaluating the contribution of genetic and environmentally-dependent variation, and in particular how these two sources of variation interact with each other, towards sexual differences. In the context of sexual size dimorphism, it has been shown that the insulin branch of the Insulin/TOR pathway enables female-specific response to nutrition. Specifically, Dilp2 mediates female-specific increase in size in the context of a protein-rich diet (Millington et al., 2021). Here we show there are several differences between males and females when comparing nutritional conditions at the level of: (1) change in mean eye size across conditions following perturbation of each Insulin/TOR component (Fig. 2C, D, Fig. 3); (2) slopes and intercepts (Fig. 4, Fig. 5); and (3) eye size variation (Fig. 5, Fig. S2). In the context of eye mean size reduction *S6k*, *Thor*, *dTOR*, *Pten*, *Tsc2* exhibited an interaction between sex and diet such that males have a greater increase while for *chico* the reverse occurred such that females have a greater increase (Fig. 3). This sex-limited reduction may be due to an perturbation-induced sexually dimorphic hypersensitivity to starved conditions as compared to wildtype. In the context of allometry, for *Pi3k* when comparing males and females across nutritional conditions, there was an interaction between sex and diet such that the eye-to-body slope decreased in females when comparing fed to starved, but it increased in males (Fig. 4 C,D,G,H Table S2). In the context of variance, for *Tsc1* when comparing males and females across nutritional conditions, there was an interaction between sex and diet such that males increased their variation while females stayed the same (Fig. S2). When looking across all three parameters (mean size, allometry, and variance), perturbation of *Pten*, the suppressor if the insulin branch of the Insulin/TOR pathway, exhibits a mosaic of sex-limited patterns. *Pten* RNAi led to double the reduction of mean eye size in males as compared to females (Fig. 2, Fig. 3), a slope increase in males like the wildtype and a slope increase in females unlike the wildtype (Fig. 5C,Table S1), and an increase in variance in males only (Table S1, Fig. S2). When considering female eye allometry in the context of perturbing *Pi3k* (female slope decreases) and *Pten* (female slope increases) together, they represent nodes capable of opposing allometric effects. This is consistent with the fact that PTEN specifically acts as a brake (dephosphorylation) for PIP2 to PIP3 conversion, antagonizing the role of PI3K, disabling AKT downstream (Georgescu, 2010). Altogether, the response of a developing male or female to nutritional plasticity in the form of changes in eye size, allometry, and variance can be similar in direction, can be limited to one sex, or the change can go in opposing directions.

### Concluding Remarks

Few complex traits have evolved across the animal kingdom to such a degree of diversity as that of the eyes (Land and Nilsson, 2012). While the evolution of the eye has baffled evolutionary biologists since Darwin, the independent origin of eyes across animals has been facilitated by an ancient eye GRN with *Pax-6* (and its paralogs) at its epicenter (Gehring and Ikeo, 1999). While the developmental basis of eye patterning and growth has been extensively studied in the model organism *Drosophila melanogaster*, how these GRNs and cell signalling pathways are impacted by nutritional plasticity in similar and different ways within and between sexes has been unexplored. Here we leverage the fly’s-eye view to illuminate how the well characterized and ancient Insulin/TOR nutrient-sensing pathway can modulate nutritional plasticity in complex ways within and between sexes through its individual pathway components. Whether these components change in function due to allelic (genetic) variation or epiallelic (epigenetic) variation to facilitate the evolution of GxE interactions should be further investigated. This work may have more general implications (Fig. 6) for sizing, size variation, trait-to-body of other organs and appendages within and between the sexes to help us understand the various molecular options to potentially facilitate morphological evolution through nutritional plasticity.

## METHODS

### *Drosophila* Stocks and husbandry

The following 15 stocks from Bloomington Drosophila Stock Centre (BDSC) were used in this study, and as described in the text, have been extensively validated: Oregon-R (5), GMR-Gal4 (1104), UAS.Empty (control; 36304), UAS.RNAi-*InR* (51518), UAS.RNAi-*chico* (36665), UAS.RNAi-*Pi3K92E* (61182), UAS.RNAi-*Akt* (33615), UAS.RNAi-*foxo (32993),* UAS.RNAi*-Tsc1* (25931), UAS.RNAi-Tsc2 (34737), UAS.RNAi-*Rheb (*33966), UAS.RNAi-*dTOR* (34639), UAS.RNAi-*raptor* (34814), UAS.RNAi-*Thor* (80427), UAS.RNAi-*S6k* (41702), UAS.RNAi-*Pten* (33643), UAS.RNAi-*impL2* (64936), UAS.RNAi-*ilp2* (32475). All flies were kept in incubators set at 25°C with a 12-hour light/dark cycle. Standard food for flies consisted of water, agar, yeast, cornmeal, sugar, propanoic acid.

### Nutritional experiments

To knockdown the Insulin/TOR pathway within the eye, an eye-specific Gal4 was crossed with an Insulin/TOR pathway component UAS-RNAi. GMR-Gal4 virgin females were crossed with males of the corresponding UAS-RNAi lines in tubes with fresh food. After 120 hours, pre-wandering 3^rd^ instar larvae from these tubes were removed from the food and gently washed. 20 individual larvae were then placed in a fed (standard food) or starved (0.8% agar-water solution) nutritional environment and left to reach the pupal stage where pupal rate and mortality did not differ between crosses. After this, individual pupae in both conditions were then moved to a new fresh food and left to reach adulthood. Wildtype Oregon-R (OrR) nutritional experiments were performed in the same way without the Gal4-UAS crosses. This protocol was adapted from starvation protocols in (Brown et al., 2019) and (Hertenstein et al., 2021).

### Microscopy (Imaging and measuring)

Adult flies that underwent nutritional conditions were sexed and imaged for measurement of their thorax length (as a proxy for body size), eye width, and eye surface area. Images were taken of *Drosophila* from head-on view using a Zeiss Axio Zoom V.16 light microscope at 29.5x magnification and measurements were taken using ZEN Pro 3.2 Software. Eye width and thorax length were measured by measuring the widest portion of the eye and longest portion of the thorax respectively. Eye surface area was measured using the surface area tool in the ZEN software and using head landmarks as seen in (Posnien et al., 2012). Averages between the left and right eye were taken for eye width and eye surface area.

### Statistics

All statistical analyses were performed on GraphPad Prism. To compare mean eye size across nutritional conditions in males and females, two-way ANOVA’s were run on log transformed data followed by Holm-Bonferroni correction for multiple comparisons. To test for changes in eye-to-body allometry within sex simple linear regressions were performed for each nutritional condition within each sex and ANCOVAs were performed to detect an effect on nutritional condition on the slope within each sex. Thorax length and eye width were plotted on XY scatter plots after being transformed using Y=log(Y)/X=log(X). Two-way ANCOVAs were preformed to detect sex*nutrition interactions for slope and y-intercept. Differences in eye size variation, while taking into account nutritional condition and sex, were assessed using a two-way Levene’s Test based on the absolute deviations of log-transformed eye width from the group median. This further enabled us to detect any interaction between sex*nutrition on variance. To determine the relative rank of the changes in variance across genotypes for fed females, starved females, fed males, starved we quantified the coefficient of variance (CV), calculated as the standard deviation divided by the mean for each group. Statistical significance: *P*<0.05

## Acknowledgments

We thank Rajakumar Lab members for both comments and both Rajakumar lab members and the Rundle lab for fly husbandry support.

## Data and resource availability

All relevant data and details of resources can be found within the article and its supplementary information

---

**Supplementary Fig. 1.**
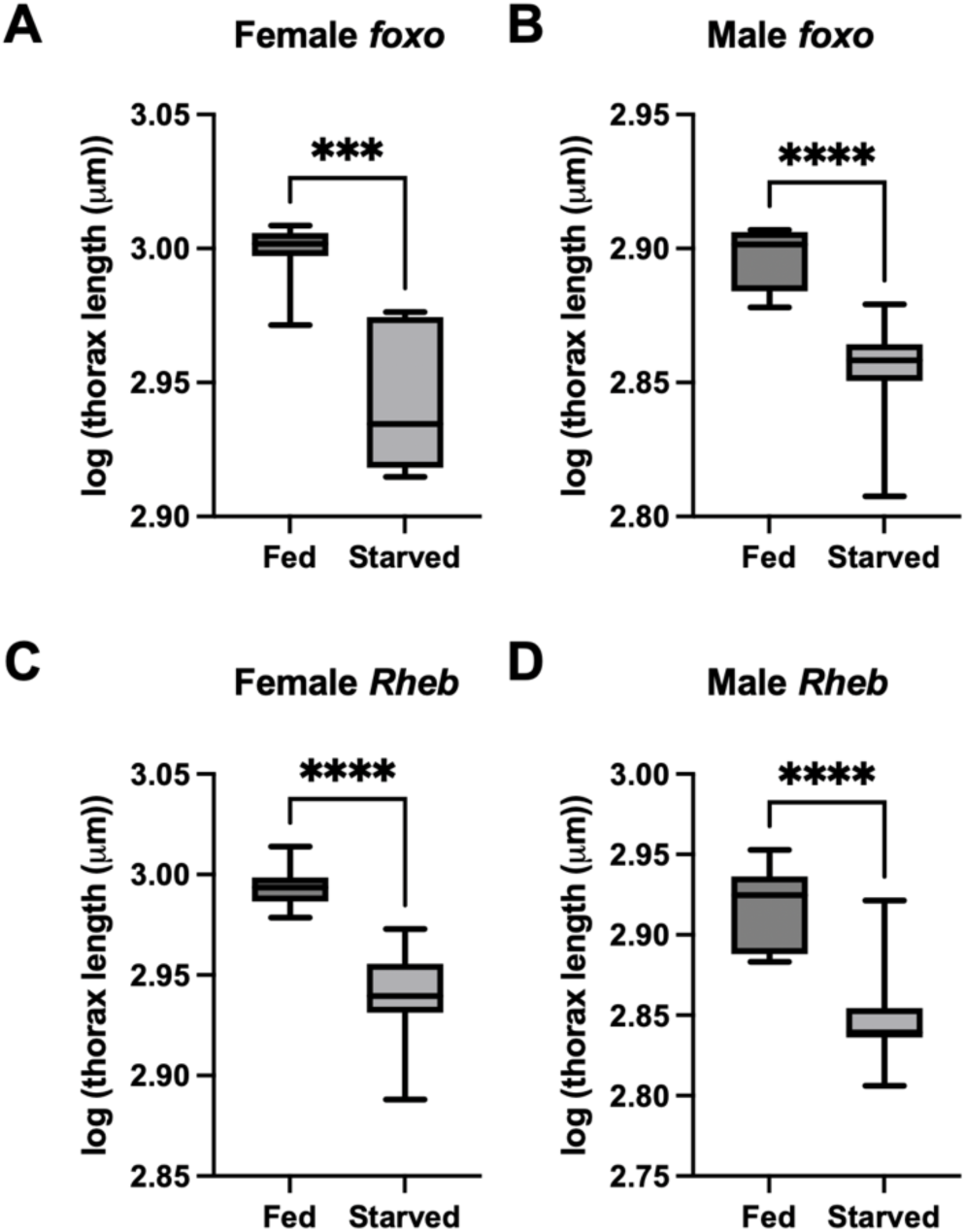
Body size comparison of fed and starved males and females with eye-specific *foxo* or *Rheb* knockdown. Thorax length comparison between fed and starved (A) females and (B) males when *foxo is* knocked down in the eye. Thorax length comparison between fed and starved (C) females and (D) males when *Rheb is* knocked down in the eye. Box of boxplots is defined by the 25^th^ and 75^th^ percentile with line at the median. Whiskers indicate min and max points. Statistical significance for Student’s *t*-tests: \*\*\**P*<0.001, \*\*\*\**P*<0.0001

**Supplementary Fig 2.**
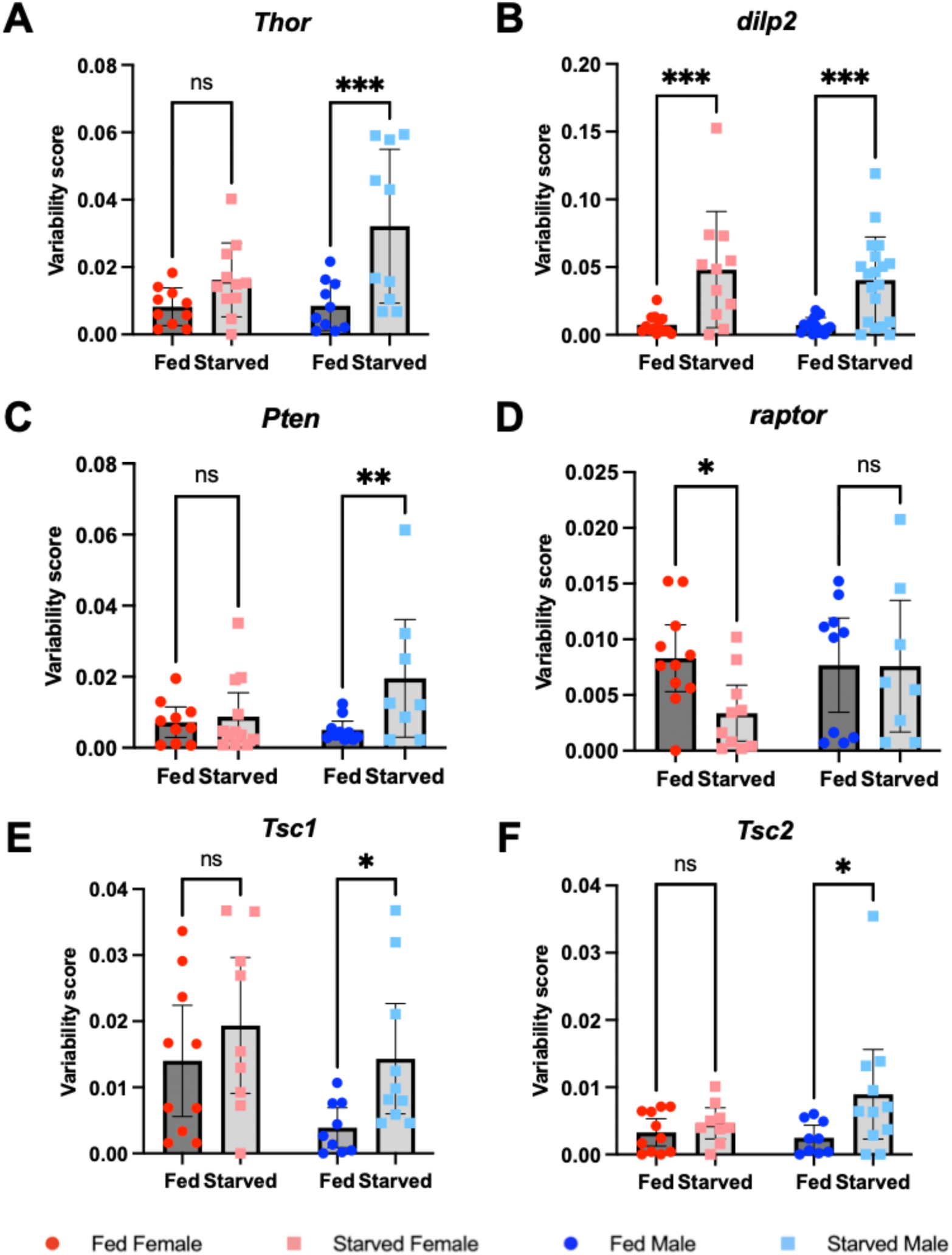
Sex- and diet-dependent variation on eye size in fed and starved Insulin/TOR knockdown females and males. Absolute deviation from the median log-transformed eye width of fed and starved males and females when (A) *Thor*, (B) *dilp2*, (C) *Pten*, (D) *raptor*, (E), *Tsc1*, or (F) *Tsc2* are knocked down in the eye (means ± 95% CI). Two-way Levene’s Test ANOVA’s were used to test for effects of sex, diet, and their interaction on variability: \**P*<0.05, \*\**P*<0.01, \*\*\**P*<0.001, ****P<0.0001. Variability score = |log(eye width) - median log(eye width)|. Female fed = dark red circles, female starved = light red squares, male fed = dark blue circles, and male starved = light blue square.

**Supplementary Table 1.** Effects of knocking down the components of the Insulin/TOR pathway on the change in slope, y-intercept, mean, and variance of eye width between fed and starved females and males. X.

| Gene | Slope |  | Y-intercept |  | Mean |  | Coefficient of Variance Rank |  |  |  |
| --- | --- | --- | --- | --- | --- | --- | --- | --- | --- | --- |
|  | Female Fed → Starved | Male Fed → Starved | Female Fed → Starved | Male Fed → Starved | Female Fed → Starved | Male Fed → Starved | Female Fed | Female Starved | Male Fed | Male Starved |
| *OrR* | No change | Increase | No change | - | Decrease | Decrease | 15 | 13 | 12 | 15 |
| Control | No change | Increase | No change | - | Decrease | Decrease | 6 | 4 | 15 | 7 |
| *InR* | No change | No change | Decrease | Increase | Decrease | Decrease | 14 | 12 | 4 | 8 |
| *chico* | No change | No change | Decrease | No change | Decrease | Decrease | 12 | 9 | 2 | 11 |
| *Pi3K* | Decrease | Increase | - | - | Decrease | Decrease | 5 | 7 | 5 | 3 |
| *Akt* | No change | No change | No change | Decrease | Decrease | Decrease | 3 | 5 | 11 | 9 |
| *Rheb* | No change | No change | No change | No change | No change | No change | 13 | 17 | 10 | 5 |
| *dTOR* | Decrease | No change | - | Increase | Decrease | Decrease | 16 | 14 | 16 | 16 |
| *raptor* | Decrease | Decrease | - | - | Decrease | Decrease | 9 | 16 | 8 | 14 |
| *S6k* | No change | No change | No change | No change | Decrease | Decrease | 4 | 8 | 1 | 10 |
| *Pten* | Increase | Increase | - | - | Decrease | Decrease | 10 | 10 | 13 | 6 |
| *foxo* | No change | No change | Increase | No change | No change | No change | 11 | 11 | 7 | 17 |
| *Tsc1* | No change | No change | No change | No change | Decrease | Decrease | 2 | 3 | 15 | 12 |
| *Tsc2* | No change | No change | No change | Decrease | Decrease | Decrease | 17 | 15 | 17 | 13 |
| *Thor* | No change | No change | Decrease | Increase | Decrease | Decrease | 7 | 6 | 6 | 2 |
| *ImpL2* | No change | No change | No change | No change | Decrease | Decrease | 1 | 2 | 3 | 4 |
| *dilp2* | No change | Increase | No change | - | Decrease | Decrease | 8 | 1 | 9 | 1 |

**Supplementary Table 2.** Changes in slopes from log-log plots of eye width-to-body allometry for Insulin/TOR pathway knockdowns in fed and starved females and males and samples size across genotypes and conditions in brackets.

| Gene | Female Fed Slope | Female Starved Slope | P-value | Male Fed Slope | Male starved Slope | P-value |
| --- | --- | --- | --- | --- | --- | --- |
| *OrR* | 0.6911 (n=27) | 0.6499 (n=24) | 0.8352 | 0.8058 (n=14) | 1.755 (n=19) | **0.0005** |
| Control | 0.6948 (n=20) | 0.3665 (n=12) | 0.4733 | 0.1364 (n=19) | 0.7570 (n=15) | **0.0053** |
| *InR* | 0.5011 (n=9) | 0.8397 (n=12) | 0.1092 | 0.1743 (n=12) | 0.1198 (n=12) | 0.8944 |
| *chico* | 0.2482 (n=9) | 0.3179 (n=12) | 0.8491 | 1.278 (n=10) | 0.5666 (n=8) | 0.0602 |
| *Pi3K* | 1.091 (n=13) | 0.2611 (n=12) | **0.0337** | -0.03234 (n=12) | 0.6421 (n=13) | **0.0069** |
| *Akt* | 0.7943 (n=10) | 0.5574 (n=13) | 0.8333 | -0.05039 (n=10) | 0.6299 (n=9) | 0.0915 |
| *Rheb* | 0.2763 (n=22) | 0.02738 (n=13) | 0.0744 | 0.1837 (n=14) | 0.6291 (n=12) | 0.1546 |
| *dTOR* | 1.448 (n=10) | 0.4517 (n=12) | **0.0041** | 0.7023 (n=10) | 0.5156 (n=8) | 0.4818 |
| *raptor* | 1.902 (n=11) | 0.5262 (n=10) | **<0.0001** | 3.27 (n=10) | 2.259 (n=8) | **0.0428** |
| *S6k* | 0.5528 (n=12) | 0.3800 (n=15) | 0.5885 | 0.5640 (n=10) | 0.6686 (n=10) | 0.6677 |
| *Pten* | -0.2007 (n=10) | 0.3197 (n=12) | **0.0178** | 0.3216 (n=10) | 1.101 (n=8) | **0.0281** |
| *foxo* | 0.4613 (n=10) | 0.3960 (n=9) | 0.7459 | 0.1242 (n=9) | 0.05794 (n=10) | 0.8402 |
| *Tsc1* | 0.6707 (n=11) | 0.8535 (n=9) | 0.8605 | 0.9677 (n=9) | 0.6165 (n=10) | 0.7330 |
| *Tsc2* | 0.4531 (n=11) | 0.8849 (n=9) | 0.1522 | 0.6733 (n=9) | 0.7065 (n=11) | 0.9371 |
| *Thor* | 0.4164 (n=10) | 0.4574 (n=11) | 0.8235 | 1.106 (n=10) | 0.7015 (n=10) | 0.4911 |
| *ImpL2* | 0.6691 (n=15) | 0.8327 (n=12) | 0.8030 | 0.4434 (n=13) | 0.9138 (n=14) | 0.4542 |
| *dilp2* | 0.2463 (n=14) | 1.045 (n=11) | 0.0710 | 0.2596 (n=16) | 0.9473 (n=19) | **<0.0001** |

Note: Bold *P*-values statistical significance ( $P < 0.05$ )

**Supplementary Table 3.** Changes in y-intercepts from log-log plots of eye width-to-body allometry for Insulin/TOR pathway knockdowns in fed and starved females and males and samples size across genotypes and conditions in brackets.

| Gene | Female Fed y-intercept | Female Starved y-intercept | P-value | Male Fed y-intercept | Male starved y-intercept | P-value |
| --- | --- | --- | --- | --- | --- | --- |
| *OrR* | 0.2889 (n=27) | 0.4085 (n=24) | 0.3604 | -0.04952 (n=14) | -2.809 (n=19) | - |
| Control | 0.2557 (n=20) | 1.222 (n=12) | 0.3702 | 1.914 (n=19) | 0.09139 (n=15) | - |
| *InR* | 0.8718 (n=9) | -0.1661 (n=12) | **<0.0001** | 1.780 (n=12) | 1.890 (n=12) | **0.0052** |
| *chico* | 1.572 (n=9) | 1.310 (n=12) | **<0.0001** | -1.437 (n=10) | 0.608 (n=8) | 0.8981 |
| *Pi3K* | -0.8953 (n=13) | 1.544 (n=12) | - | 2.404 (n=12) | 0.4141 (n=13) | - |
| *Akt* | -0.06256 (n=10) | 0.6272 (n=13) | 0.2069 | 2.428 (n=10) | 0.4120 (n=9) | **0.0081** |
| *Rheb* | 1.524 (n=22) | 2.267 (n=13) | 0.7962 | 1.772 (n=14) | 0.4957 (n=12) | 0.4895 |
| *dTOR* | -1.961 (n=10) | 0.9912 (n=12) | - | 0.2825 (n=10) | 0.7949 (n=8) | **<0.0001** |
| *raptor* | -3.276 (n=11) | 0.7949 (n=10) | - | -7.222 (n=10) | -4.222 (n=8) | - |
| *S6k* | 0.6764 (n=12) | 1.18 (n=15) | 0.3240 | 0.6554 (n=10) | 0.3399 (n=10) | 0.0907 |
| *Pten* | 2.979 (n=10) | 1.418 (n=12) | - | 1.423 (n=10) | -0.8146 (n=8) | - |
| *foxo* | 0.9802 (n=10) | 1.195 (n=9) | **0.0004** | 1.95 (n=9) | 2.139 (n=10) | 0.7013 |
| *Tsc1* | 0.3569 (n=11) | -0.1860 (n=9) | 0.7037 | -0.5230 (n=9) | 0.5200 (n=10) | 0.5806 |
| *Tsc2* | 1.03 (n=11) | -0.2434 (n=9) | 0.1097 | 0.3767 (n=9) | 0.2568 (n=11) | **0.0002** |
| *Thor* | 1.133 (n=10) | 0.9738 (n=11) | **<0.0001** | -0.871 (n=10) | 0.2426 (n=10) | **<0.0001** |
| *ImpL2* | 0.3375 (n=15) | -0.1523 (n=12) | 0.8942 | 1.017 (n=13) | -0.3490 (n=14) | 0.7082 |
| *dilp2* | 1.610 (n=14) | -0.7681 (n=11) | 0.2569 | 1.560 (n=16) | -0.4527 (n=19) | - |

Note: Bold *P*-values statistical significance ( $P<0.05$ )

